## Supplementary Material for "Forecasting protein evolution by integrating birth-death population models with structurally constrained substitution models"

The supplementary material includes Tables S1-S3, Figure S1 and references.

### Supplementary tables

**Table S1. Evolutionary processes and corresponding parameters implemented in *ProteinEvolver2*.** The user can specify a variety of parameters, optional or mandatory, to define an evolutionary scenario.

| Evolutionary process | Parameter | Mandatory or optional \| | *Additional information* |
|---|---|---|---|
| All | Number of replicates | M | |
| Evolutionary history | Birth-death process (includes parameters presented below) | | *One of these options must be used* |
| Evolutionary history | Coalescent process (includes parameters presented below) | M | |
| Evolutionary history | Input phylogenetic tree/s (fixed tree in *Newick* format) | | |
| DNA evolution / Substitution model of DNA evolution | Nucleotide frequencies | O | *By default, the JC model is applied* |
| DNA evolution / Substitution model of DNA evolution | Transition / transversion ratio | O | |
| DNA evolution / Substitution model of DNA evolution | Relative symmetrical substitution rates | O | |
| DNA evolution / Substitution model of DNA evolution | Relative asymmetrical substitution rates | O | |
| DNA evolution / Substitution model of DNA evolution | SCS models[1] for DNA | O | |
| Protein evolution / Substitution models of protein evolution | Empirical amino acid substitution model (it implements a variety of empirical models)[1] | | *One of these options must be used* |
| Protein evolution / Substitution models of protein evolution | SCS models[2] for proteins | M | |
| Molecular evolution / Substitution models of evolution | Rate variation among sites[3] | O | |
| Molecular evolution / Substitution models of evolution | Proportion of invariable sites | O | |
| Molecular evolution / Substitution rate | Variable site-specific substitution rate | O | |
| Molecular evolution | User-specified sequence for the root node | O[4] | |
| Information in output files | Print sequences to a file | O | |
| Information in output files | Format of simulated multiple sequence alignments (*Fasta, Phylip, Nexus*) | O | *By default, printed in Phylip format* |
| Information in output files | Print sequence of the root node (GMRCA or MRCAs) | O | |
| Information in output files | Print simulated trees | O | |
| Information in output files | Print times of nodes of genealogies | O | |
| Information in output files | Print simulated ARG | O | |
| Information in output files | Print recombination breakpoints | O | |
| All | Simulation seed | O | |
| Information in the screen | Level of information printed on the screen | O | |
| Evolutionary history / Birth-death evolutionary process | Type of birth and death rates (specified or calculated) | M | *Birth and death rates* |

| | | | |
|---|---|---|---|
| | | | *are specified or calculated from the fitness of the variant* |
| Global birth-death rate variation among lineages following (Neher *et al*, 2014) | Model option for scenarios based on fitness, where death rate is 1 and birth rate is 1 + fitness | O | *Requires indicating No (1) or Yes (1)* |
| Evolutionary history / Birth-death evolutionary process | Type of ending the simulation of the birth-death process | M | *Reaching a specified sample size, number of tip nodes or evolutionary time* |
| Evolutionary history / Birth-death evolutionary process | Prune extinct nodes | O | |
| Evolutionary history / Birth-death evolutionary process | Outgroup and its branch length | O | |
| Evolutionary history / Birth-death evolutionary process | Substitution rate | M | |
| Evolutionary history / Birth-death evolutionary process | Effective population size; Haploid/Diploid | M | |
| Evolutionary history / Birth-death evolutionary process | Alignment length (nucleotides or amino acids) | M | |
| Evolutionary history / Coalescent evolutionary process | Sample size and alignment length (nt or aa) | M | |
| Evolutionary history / Coalescent evolutionary process | Effective population size; Haploid/Diploid | M | |
| Evolutionary history / Coalescent evolutionary process | Tip dates[5] | O | |
| Evolutionary history / Coalescent evolutionary process | Generation Time | O | |
| Evolutionary history / Coalescent evolutionary process | Exponential growth rate | O | |
| Evolutionary history / Coalescent evolutionary process | Demographic periods | O | |
| Evolutionary history / Coalescent evolutionary process | Migration model (island, stepping-stone, island-continent) and population structure | O | |
| Evolutionary history / Coalescent evolutionary process | Migration rate (constant or variable with time) | O | |
| Evolutionary history / Coalescent evolutionary process | Convergence of demes | O | |
| Evolutionary history / Coalescent evolutionary process | Homogeneous recombination rate | O | |
| Evolutionary history / Coalescent evolutionary process | Fixed number of recombination events | O | |
| Evolutionary history / Coalescent evolutionary process | Recombination hotspots | O | |
| Evolutionary history / Coalescent evolutionary process | Substitution rate | M | |
| Evolutionary history / Coalescent evolutionary process | Outgroup and its branch length | O | |

| | | |
|---|---|---|
| Evolutionary history / User-specified phylogenetic tree/s | Number of input phylogenetic trees, alignment length (in nucleotides or amino acids) and rooted phylogenetic tree | M, M, M |
| Molecular evolution | Sequence length | M |
| Molecular evolution | Factor that multiplies the original substitution rate | O |
| Molecular evolution | Sequenced assigned to the root node | O |
| Molecular evolution / Structurally constrained substitution models of protein evolution | PDB file | M |
| Molecular evolution / Structurally constrained substitution models of protein evolution | Chain of the PDB file | M |
| Molecular evolution / Structurally constrained substitution models of protein evolution | Input file of amino acid contacts | M |
| Molecular evolution / Structurally constrained substitution models of protein evolution | Thermodynamic temperature | M |
| Molecular evolution / Structurally constrained substitution models of protein evolution | Configurational entropy per residue (unfolded) | M |
| Substitution models of protein evolution | Configurational entropy per residue (misfolded) | M |
| Molecular evolution / Structurally constrained substitution models of protein evolution | Configurational entropy offset (misfolded) | M |
| Molecular evolution / Structurally constrained substitution models of protein evolution | Third cumulant in REM calculation | M |
| Molecular evolution / Structurally constrained substitution models of protein evolution | Type of SCS model (Neutral or Fitness) | M |
| Molecular evolution / Structurally constrained substitution models of protein evolution | Effective population size for the fitness SCS model | O |
| Molecular evolution / Structurally constrained substitution models of protein evolution | Consideration of branch lengths | O |
| Molecular evolution / Structurally constrained substitution models of protein evolution | Amount of information about SCS models printed as output files | O |

[1]A variety of empirical substitution models of protein evolution are implemented: *Blosum62* (Eddy, 2004; Henikoff & Henikoff, 1992), *CpRev* (Adachi *et al*, 2000), *Dayhoff* (Dayhoff *et al*, 1978), *DayhoffDCMUT* (Kosiol & Goldman, 2005), *FLU* (Dang *et al*, 2010), *HIVb* (Nickle *et al*, 2007), *HIVw* (Nickle *et al.*, 2007), *JTT* (Jones *et al*, 1992), *JonesDCMUT* (Kosiol & Goldman, 2005), *LG* (Le & Gascuel, 2008), *Mtart* (Abascal *et al*, 2007), *Mtmam* (Yang *et al*, 1998), *Mtrev24* (Adachi & Hasegawa, 1996), *RtRev* (Dimmic *et al*, 2002), *VT* (Muller & Vingron, 2000), *WAG* (Whelan & Goldman, 2001) or any user-specified matrix for all the sites or for every site (thus differing among sites).

[2]SCS models can be neutral or fitness-based landscape.

[3]Shape of the gamma distribution.

[4]If not specified, a random sequence is assigned to the root node according to the used nucleotide, codon or amino acid frequencies.

[5]In presence of convergence of demes, the tip nodes must be older than the convergence of demes.

**Table S2. Longitudinal data of the HIV-1 PR used to evaluate the accuracy of the forecasting protein evolution.** For each patient, the first column indicates the identifier code (ID) of the patient in the Specialized Assistance Services in Sexually Transmissible Diseases and HIV/AIDS in Brazil. The next columns indicate, for every consensus sequence collected at a time *T* from the patient, the GenBank accession code and the number of amino acid substitutions accumulated since *T1* (shown in parenthesis). The last column indicates the HIV-1 PR inhibitor/s that the patient received.

| Patient ID | **Longitudinal sample (*T*)** | | | | | HIV-1 PR inhibitors administrated |
|---|---|---|---|---|---|---|
| | ***T1 origin (# of substitutions)*** | ***T2 (# of substitutions)*** | ***T3 (# of substitutions)*** | ***T4 (# of substitutions)*** | ***T5 (# of substitutions)*** | |
| 99842856 | ON983124 (0) | ON983123 (3) | ON983126 (9) | ON983124 (11) | - | RTV, LPV |
| 99943945 | ON982892 (0) | ON982893 (5) | ON982894 (8) | ON982891 (11) | - | RTV, FPV |
| 10701966 | ON982841 (0) | ON982837 (6) | ON982838 (12) | ON982839 (17) | ON982840 (19) | ATV, RTV, LPV |
| 12887 | ON982884 (0) | ON982883 (3) | ON982885 (6) | ON982886 (8) | - | DRV, RTV |
| 99817844 | ON982842 (0) | ON982843 (5) | ON982845 (9) | ON982844 (12) | - | ATV, RTV |
| 99654931 | ON983078 (0) | ON983079 (10) | ON983077 (19) | ON983110 (22) | - | LPV |
| 99574196 | ON982995 (0) | ON983050 (10) | ON982996 (17) | ON982994 (19) | - | LPV |
| 37000881 | ON983034 (0) | ON983035 (1) | ON983036 (6) | ON983033 (8) | - | ATV, RTV, LPV |
| 99412571 | ON983119 (0) | ON983120 (3) | ON983121 (5) | ON983122 (7) | - | - |
| 99783386 | ON982927 (0) | ON982925 (4) | ON982924 (9) | ON982928 (12) | ON982926 (17) | LPV, RTV, ATV |
| 23400093 | ON983128 (0) | ON983130 (12) | ON983129 (18) | ON983127 (22) | - | DRV, RTV, ATV, LPV |
| 4300388 | ON982876 (0) | ON982875 (5) | ON982877 (11) | ON982878 (20) | - | LPV, DRV, RTV |

**Table S3. Structures of the HIV-1 MA, SARS-CoV-2 Mpro, Influenza NS1 protein and, SARS-CoV-2 PLpro. Also structures of the HIV-1 PR selected as templates in homology modeling.** The data of the HIV-1 matrix (MA) protein, influenza NS1 protein, and SARS-CoV-2 main protease (Mpro) and papain-like protease (PLpro) involved only one consensus sequence at the initial time, thus only one protein structure was used and did not require homology modelling because the study sequence was already present in an available protein structure. However, the HIV-1 protease (PR) dataset involved several independent HIV-1 populations (patients) and, a structural template was selected for each one. For each protein, the table shows the corresponding PDB code and protein chain used for the study. For the HIV-1 PR, the table presents the modelling quality and sequence identity of the structural templates for homology modelling (their coverage was 100%).

| **HIV-1 MA** | |
|---|---|
| *PDB code* | *Protein chain* |
| 7JXR | B |
| **SARS-CoV-2 Mpro** | |
| *PDB code* | *Protein chain* |
| 7N8C | A |
| **SARS-CoV-2 PLpro** | |
| *PDB code* | *Protein chain* |
| 6XA9 | A |
| Influenza NS1 | |
| *PDB code* | *Protein chain* |
| 4OPH | A |

| **HIV-1 PR** | | | | |
|---|---|---|---|---|
| *Patient ID* | *PDB code* | *Protein chain* | *Modelling quality* | *Sequence identity* |
| 99842856 | 3LZS | A | 555.3381 | 88.889 |
| 99943945 | 1C6X | A | 453.8610 | 83.838 |
| 10701966 | 3U71 | A | 467.8960 | 95.960 |
| 12887 | 3EKP | A | 447.4938 | 80.808 |
| 99817844 | 1HIV | A | 641.8560 | 90.816 |
| 99654931 | 3LZS | A | 464.9483 | 90.909 |
| 99574196 | 1AID | A | 632.7499 | 93.939 |
| 37000881 | 1AID | A | 544.9096 | 92.929 |

| | | | | |
|---|---|---|---|---|
| 99412571 | 3D3T | A | 529.7580 | 90.909 |
| 99783386 | 2FDE | A | 450.4926 | 94.949 |
| 23400093 | 1SGU | A | 435.4226 | 91.818 |
| 4300388 | 2FDD | E | 638.4982 | 83.838 |

### Supplementary figures

**Figure S1. Distribution of amino acid substitutions observed along the HIV-1 MA sequences at time *T31*.** Left: Distribution of the observed amino acid substitutions along the HIV-1 matrix (MA) protein sequences at time *T31*. Right: Distribution of the indicated amino acid substitutions (shown in blue) along the protein structure.

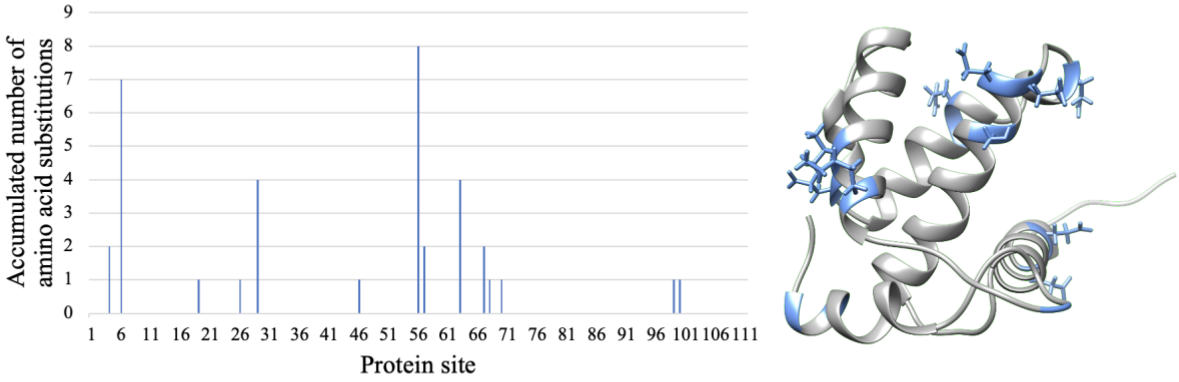
